# Reprogrammed *Pteropus* Bat Stem Cells Present Distinct Immune Signature and are Highly Permissive for Henipaviruses

**DOI:** 10.1101/846410

**Authors:** Noémie Aurine, Camille Baquerre, Maria Gaudino, Christian Jean, Claire Dumont, Sylvie Rival-Gervier, Clémence Kress, Branka Horvat, Bertrand Pain

**Affiliations:** University of Lyon, Université Lyon 1, INSERM, INRA, Stem Cell and Brain Research Institute, U1208, USC1361, Bron, France; CIRI, International Center for Infectiology Research, University of Lyon, Université Claude Bernard Lyon 1, INSERM, U1111, CNRS, UMR5308, ENS Lyon, Lyon, France

**Keywords:** Bats, Stem cells, Reprogramming, Emerging infection, Nipah virus, Innate immunity, transcriptomic analysis, interferon-stimulated gene

## Abstract

Bats are unique among mammals due to the ability of powered flight and exceptional longevity. They are also asymptomatic hosts for numerous viruses, including recently emerged zoonotic Henipaviruses Nipah and Hendra, which are highly pathogenic for humans and other mammals. Better understanding of how bats control viral infection requires development of relevant permissive cellular experimental models. By applying a somatic reprogramming protocol to *Pteropus* bat primary cells, using a novel combination of ESRRB, CDX2, and c-MYC transcription factors, we generated bat reprogrammed cells exhibiting stem cell-like characteristics and a neural stem cell-like molecular signature. These cells present a unique interferon-stimulated transcriptomic signature and both produce and respond to interferon type-I, highlighting differences between stem cells from bats and other mammals. In contrast to primary bat cells, these reprogrammed cells are highly susceptible to infection by *Henipavirus*, thereby enabling isolation of new bat viruses, study of virus-bat interactions, and better understanding of bat biology.

**Summary sentence:** Somatic reprogramming provides new bat stem cells with unique immune properties and original viral permissivness

## Introduction

Emergence of zoonotic diseases is increasing globally; thus the ability to predict and prevent viral epidemics is a major objective for public health organizations. Among all emerging infectious diseases, approximately 60% are of zoonotic origin^1^. Bats host a large number of human and animal zoonotic viruses, including Henipaviruses^2^ (Nipah (NiV) and Hendra (HeV)), Coronaviruses (Severe Acute Respiratory Syndrome (SARS-CoV)^3^ and Middle East Respiratory (MERS-CoV))^4^, and Filoviruses (Marburg^5^ and Ebola viruses^6^). These agents are among the most virulent pathogens emerging from animal reservoirs and are capable of infecting a broad range of species, including humans, while remaining asymptomatic in bats. Transmission of NiV from bats to humans occurs either through intermediate host contamination^7,8^ or directly^9^ via consumption of fruits or raw date palm juice contaminated with bat saliva or urine. In the case of NiV-Bangladesh isolates, this may be followed by inter-human transmission^10^. Henipaviruses are negative strand RNA viruses; both NiV and HeV are responsible for outbreaks of respiratory and neurological diseases in Southeast Asia and Australia, respectively, with a fatality rate of 40–100%^11^. NiV and Henipa-like viruses have been detected molecularly and/or serologically in *Pteropus* bats from different Asian and African Countries^12^, and worldwide distribution of these bat species poses a threat of future NiV pandemics^13^.

Bats are an ancient and diverse group of animals (the second largest order of mammals after rodents), representing around 20% of all mammalian species^14^. Indeed, they are the most abundant and widely distributed non-human mammalian species worldwide^15^. Bats, the only mammals capable of powered flight, show extraordinary longevity compared with their corporal mass^16,17^. Phylogenetic analyses based on molecular data classified bats into two suborders: Yangochiroptera (microbat families) and Yinpterochiroptera (including the non-echolocating *Pteropodidae* family, which contains the *Pteropus genus* and the echolocating *Rhinolophoidea*)^18^. A large number of novel bat virus sequences was identified in these two bat suborders, which diverged over 50 million years ago, suggesting a long co-evolutionary history of bats and their viruses that have shaped unique host-pathogen relationships^19^. In addition, bats harbor a markedly higher proportion of zoonotic viruses than other mammalian orders^20,21^. Although bat serology and viral genome sequencing permit detection of a number of bat viruses, isolating them has been far more challenging. Studying the bat immune system and bat/virus interactions provides valuable insight into the mechanisms underlying successful control of viral infection, and may lead to novel approaches to managing viral spillover and development of new antiviral strategies for humans^22^. As infection studies in bats are difficult to implement due to the animal facility constraints and the absence of bat husbandry, it is crucial to establish adequate research tools for comparative *in vitro* infection, for studying virus-host interactions, and for isolating new bat viruses^23^.

Availability of relevant cell lines is particularly important and presents a critical obstacle for further study of virus-host interactions^24^. In humans, the cellular targets of Henipaviruses are endothelial, epithelial, and neural cells. Although several cell lines from *Pteropus* bats have been established, most are immortalized primary cells (PCs) with fibroblast-like morphology. Utilization of high multiplicities of infection by most studies^25^ suggests that these cells show low permissiveness for *Henipavirus* infection and replication. The use of primary and immortalized cells limits our ability to isolate and study viruses that may show tropism for cell types that are more difficult to establish as a primary culture or to immortalize, thereby emphasizing the need to develop more appropriate *Pteropus* bat cells with different phenotypes.

Pluripotent stem cells (PSCs) are able to self-renew *in vitro* and differentiate into derivatives of the three embryonic germ layers: endoderm, ectoderm, and mesoderm. Somatic reprogramming generates induced PSCs (iPSCs) via four pluripotent transcription factors, known as OSKM (a combination of the POU5F1/OCT4, SOX2, KLF4 and c-MYC genes)^26^. Initially demonstrated in mice, then in humans, rats, and non-human primates, the process has been tested (with variable success) in a large number of mammalian species, including rabbits, sheep, bovines, pigs, horses, dogs, and even in some endangered species^27–29^; however, despite the importance of bats to ecology and immunovirology, and although certain attempts have been made in that direction^30^, somatic reprogramming of primary bat cells has not been examined rigorously.

Here, we generated *Pteropus* bat reprogrammed stem cells (RSCs) from PCs using a novel combination of three transcription factors: ESRRB, CDX2 and c-MYC. These cells exhibit typical PSC morphology, are recognized by SSEA-1 and EMA-1 stem-cell-specific antibodies, display PSC-specific chromatin features, and express a set of neural stem cell-associated genes. These reprogrammed cells are much more highly susceptible to virus infection than the parental primary bat cells, making them a new cell type that can be used to isolate novel bat viruses and study host-pathogen interactions. Interestingly, in contrast to stem cells from other mammalian species, bat RSCs are capable of both responding to and producing IFN-I, and present a unique interferon-stimulated gene (ISG) transcriptomic signature, suggesting some fundamental differences between bats and other mammals. This new cell model is an important experimental tool that will improve understanding of the unique properties of bat/virus interactions and will help to decipher novel aspects of bat biology.

## Results

### *Pteropus* bat PCs show limited proliferation *in vitro*

Explants derived from *Pteropus* bat trachea (PTC), lung (PTL), and alary membrane skin (PAC) were cultured in gelatin-coated wells either in fibroblast medium (FM) or in ES derived medium (ESM1). Individual cells emerged from these explants at 5, 15, and 30 days after plating, respectively; all cells exhibited a fibroblastic morphology, with an elongated and often flattened shape (Fig. 1a–c). PLCs entered senescence after 4 to 5 generations, while PACs became senescent after 10–11 generations; by contrast, PTCs continued to proliferate for more than 30 generations (200 days of continuous culture), although there was a slow, progressive and irreversible decrease in the proliferation rate during that time (Fig. 1d). These results strongly suggested that primary cultures obtained from different types of bat tissue have a limited life span *in vitro*, similar to that of other mammalian PC cultures.

**Fig. 1.**
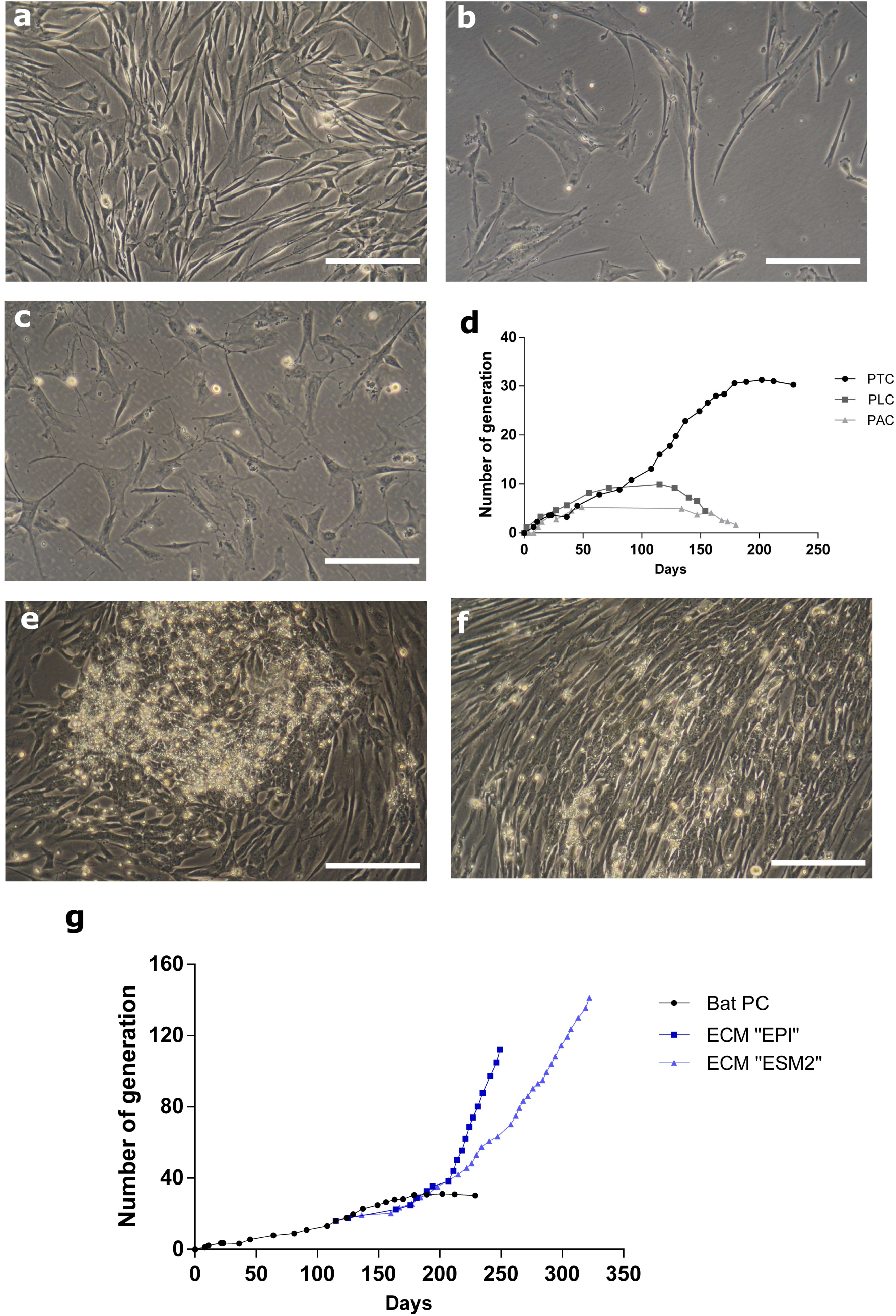
Generation of bat reprogrammed stem cells. Bat primary cell cultures (bat PC) derived from trachea (PTC) (a), lung (PLC) (b), and alary membrane (PAC) (c) observed under a light microscope. (d) Comparative growth curves of *Pteropus* primary cells. Somatic reprogramming of bat PCs with inducible transposons encoding CDX2, c-MYC, and ESRRB genes and culture in EpiStem (EPI) (e) or ESM2 (f) media. (g) Comparative growth curves of bat PC and reprogrammed bat stem cells (bat RSCs). Scale bar, 200 μm.

### Reprogramming of primary *Pteropus* bat cells

We wanted to obtain bat cells with stem cell features and the ability to proliferate long-term but did not want to use immortalizing agents. Therefore, we used a somatic reprograming approach to introduce selected “reprogramming” genes into PCs. Based on the partially annotated genome of *P. vampyrus*, alignment of protein sequences derived from human and *P. vampyrus* genes revealed identities ranging from 87–99% between seven major reprogramming human genes, with the noticeable exception of NANOG, which showed only 70% identity (Supplementary Table 1). Therefore, we first used Sendai viruses to introduce the human OSKM gene combination into bat PCs in the presence or absence of NANOG, which was expressed through a transposon. Although the genes were delivered and expressed efficiently (Supplementary Fig. 1b), no long-term morphological changes were observed up to 100 days post-infection (Supplementary Fig. 1a). Delivering the same gene combination using inducible transposon vectors did not provide a better result, although an increase in cellular proliferation was observed, along with transient and partial morphological changes (Supplementary Fig. 1c, d).

Next, we screened additional transcription factors for their ability to induce somatic reprogramming of bat PCs through delivery of doxycycline-inducible transposons and identified a novel gene combination (ESRRB, CDX2, and c-MYC) as highly efficient at inducing marked and stable morphological changes in transduced bat PCs. When cells were cultured in serum-free EPI medium, foci showing typical stem cell morphology appeared at around 25 days after addition of doxycycline (Fig. 1e); however, a longer period (35 days) was required when cells were cultured in serum-containing ESM medium (Fig. 1f). Stable proliferation of bat RSCs has continued for more than 110–140 generations, spanning 250–320 days (Fig. 1g). The average doubling time in EPI medium was estimated to be around 24 h, and that in ESM medium around 35 h.

### Bat RSCs show stem cell-like characteristics

Bat RSCs grew in compact colonies in both EPI and ESM medium, with a morphology typical of PSCs: small and round, with a large nucleus, a high nucleo-cytoplasm ratio, and a prominent nucleolus (Fig. 2a). Electron microscopy confirmed that the morphology and ultrastructure of bat RSCs was very different from that of bat PCs. In particular, chromatin was distributed homogeneously throughout the nucleoplasm, without large zones of electron-dense heterochromatin that are typical of differentiated cells, and regularly observed in the nuclei of bat PCs. The endoplasmic reticulum in bat RSCs was abundant, but not dilated as in bat PCs. (Fig. 2b). Cell cycle analysis of bat RSCs revealed a pluripotent stem cell-like profile with a short G2/M phase and a long S phase, unlike bat PCs. Expression of PSC-specific antigens, including stage-specific embryonic antigens (SSEA-1, SSEA-3, and SSEA-4) and an epithelial membrane antigen (EMA)-1^31,32^, was then analyzed by immunostaining and flow cytometry (Fig. 2d). Similar to murine ESCs (mESCs), but in contrast to human PSCs (hiPSCs), bat RSCs were positive for SSEA-1 and EMA-1 and negative for SSEA-3 and SSEA-4. The nuclear distribution of some epigenetic marks in RSCs was similar to the pattern observed in some PSCs (Fig. 2f). Indeed, the facultative heterochromatin marker H3K27me3 in bat RSCs was distributed within several large foci, rather than being concentrated mainly at the inactivated X chromosome as in PCs. Previously, such numerous prominent H3K27me3 foci were observed in both naïve mouse PSCs33 and in pluripotent and reprogrammed avian cells^31,34^. Taken together, these different expression profiles strongly suggest that following reprogramming, bat RSCs share key features with PSCs from other species.

**Fig. 2.**
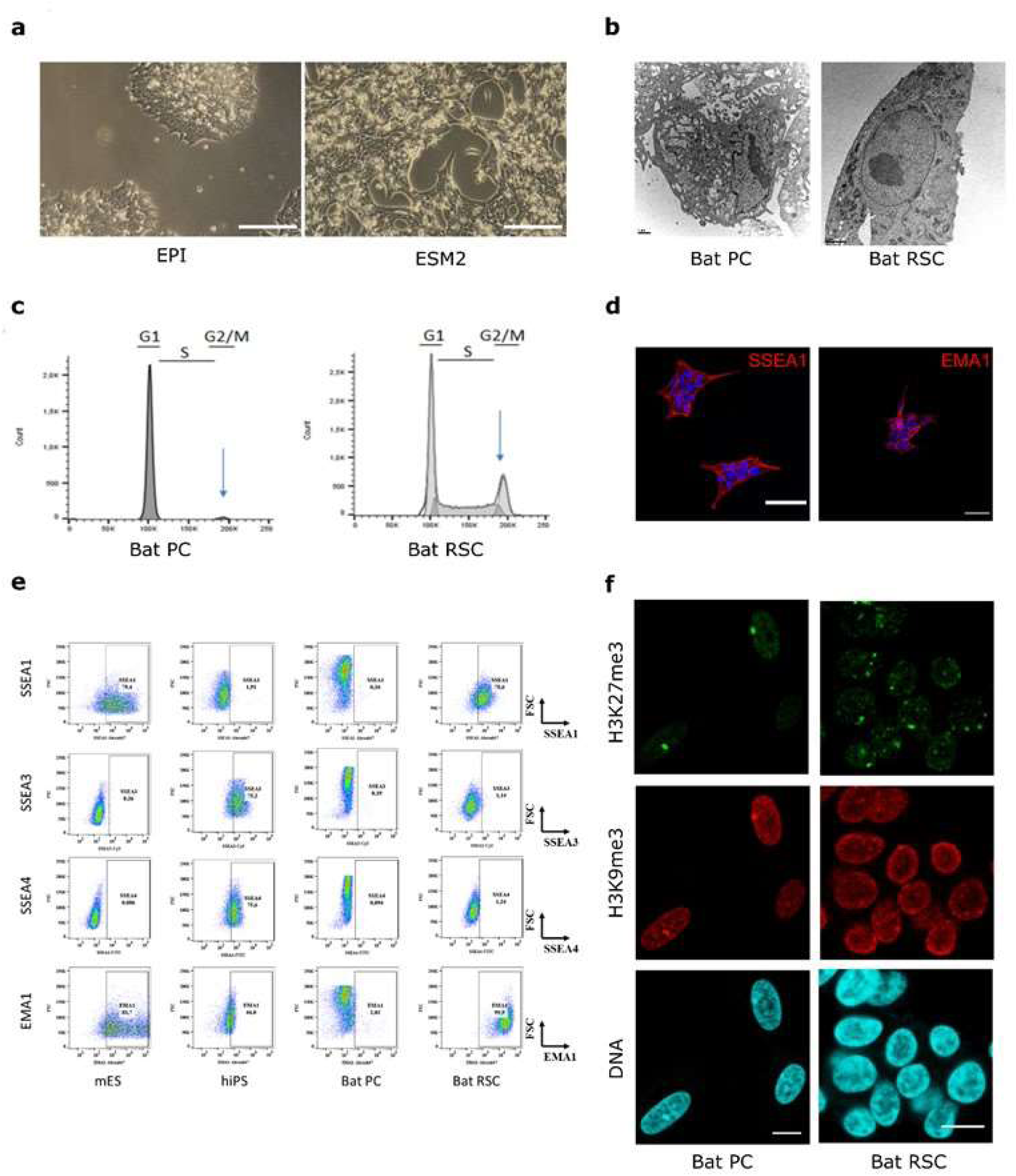
Characterization of *Pteropus* bat reprogrammed stem cells. (a) Morphology of bat RSCs after 170 days of culture in EPI or ESM2 medium. Scale bar, 200 μm. (b) Ultrastructural analysis of reprogrammed cells by electron microscopy. Scale bar, 1 μm. (c) Cell cycle analysis using propidium iodide staining. Expression of markers of pluripotency expressed by bat PCs and bat RSCs, as analyzed by immunostaining (d) and flow cytometry (e) and compared with expression by murine embryonic stem cells (mESC) and human pluripotent stem cells (hiPSC). (f) Histone post-translational modifications in reprogrammed cell nuclei: immunodetection of H3K27me3 and H3K9me3, and DNA counterstaining with TO-PRO-3. Scale bar, 10 µm.

### Bat RSC have a specific neural stem cell molecular signature

To define the full transcriptomic landscape of generated bat RSCs, we performed deep sequencing of RNA isolated from primary and reprogrammed cells. Principal component analysis (PCA) of expressed genes indicated that bat RSCs cultured in either EPI or ESM medium had expression profiles clearly distinct from those of bat PCs (Fig. 3a). Furthermore, 3506 genes were differentially expressed between bat PCs and bat RSCs, with a log -fold change > 2 (LFC > 2) and a p-value > 0.05 (Supplementary Table 2). When cultured in either medium during reprograming, bat RSCs did not express key genes related to pluripotency, such as POU5F1/OCT4 and NANOG; noticeable exceptions were SOX2 and ZSCAN4, which play roles in telomere maintenance and long-term-genomic stability in ES cells^35^. This suggests that the reprogramming factors introduced into the cells induced a new cell type. String analysis on the first 823 differentially expressed genes (DEGs) with a LFC > 5 between bat PCs and bat RSCs revealed the presence of several major gene clusters (Supplementary Table 3). Further GO analysis of these 823 DEGs indicated that the first three GO terms for biological processes were “nervous system development” (GO:0007399), “generation of neurons” (GO:0048699) and “neurogenesis” (GO:0022008) (Supplementary Table 4). The most important GO terms for molecular function were “inorganic molecular entity transmembrane transporter activity” (GO:0015318), “ion transmembrane transporter activity” (GO:0015075), and “metal ion transmembrane transporter activity” (GO:0046873). The cellular component GO terms included “plasma membrane part” (GO:0044459), “plasma membrane” (GO:0005886), and “cell periphery” (GO:0071944) (Supplementary Table 4). Additional analysis of the main gene clusters revealed that the most important cluster contained 27 members (Cluster 27, Fig. 3b) and was centered on GRIA1 and GRIK2; this cluster harbored numerous specific neural-associated receptors for neurotransmitters (e.g., GABA and 5-Hydroxytryptamine (serotonin)), illustrating the neural cell-like nature of RSCs. The second most important cluster centered on SOX2 and contained several transcription factors, some of which are specific to PSCs: POU3F3, NKX2-5, GATA4, ZIC2 and ZIC5, CDX2, MYOD1, HNF4A, PRDM14, PIWIL2, TBR1, FOXA1 and FOXA3, PITX3, LIN28B, ELAVL2, and ZSCAN4 (Cluster 23a, Fig. 3c). Other clusters contained neural markers and processes, ion channels and membrane bound transporters, and some components participating in the G protein-coupled receptor signaling pathway (Supplementary Table 3). These expression profiles suggest that the reprogramming process gave rise to a new neural-related stem cell type that differs from the initial PCs and shares features with PSCs described in other species. Interestingly, ephrin B2 (EFNB2), a receptor for *Henipavirus* entry, was among the main DEGs and was expressed at much higher levels in bat RSCs than in bat PCs (Supplementary Table 2). Such an expression profile suggests the possibility of *Henipavirus* entry into these cells. We tested this in the following experiments.

**Fig. 3.**
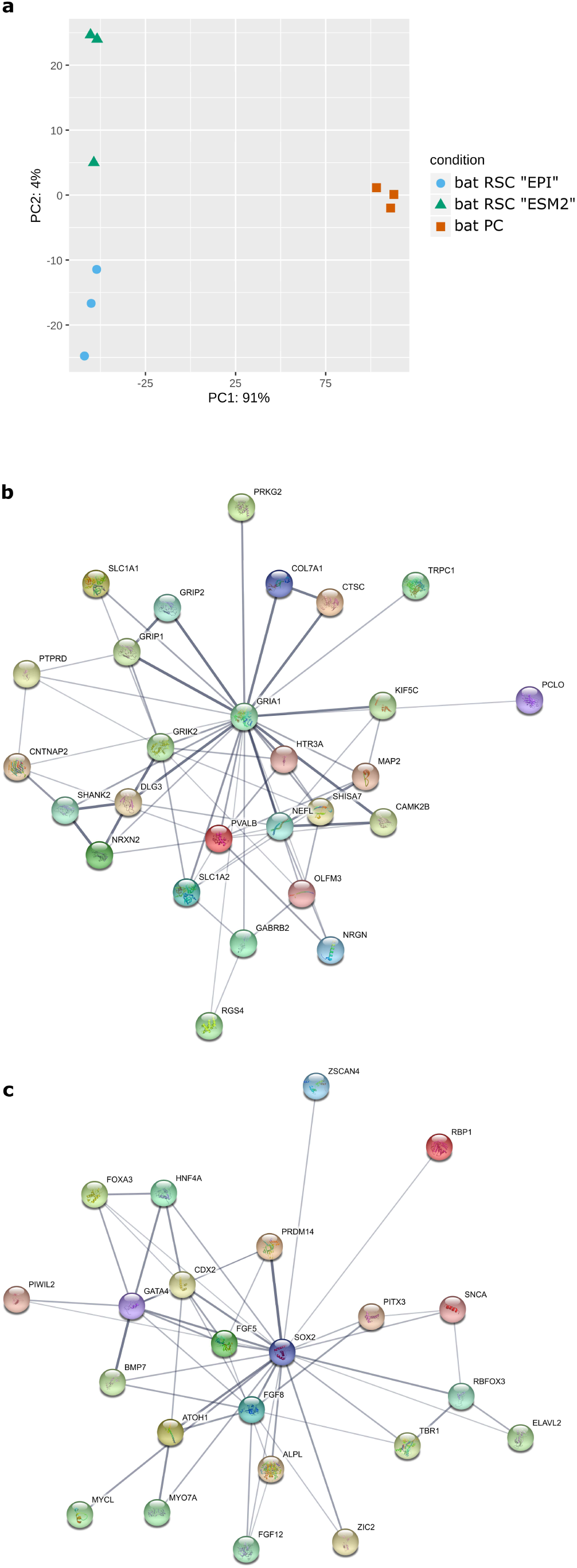
Molecular features of *Pteropus* bat reprogrammed stem cells. (a) Principal component analysis (PCA) of RNA-seq results for bat PCs and bat RSCs. Principal Component 1 (PC1, x-axis) represents 91% and PC2 (y-axis) represents 4% of the total variation in the data. (b) Protein network analysis of the first main cluster of 27 genes identified by String analysis of the DEGs identified by RNAseq analysis and centered on GRIA1 and GRIK2 genes (c) Protein network analysis of the second main cluster of 23 genes, centered on the SOX2 gene.

### Bat RSCs are highly permissive for Henipavirus infection

As *Pteropus* bats are the recognized natural reservoir of Henipaviruses, we tested the susceptibility of bat PCs and bat RSCs to infection by two human NiV isolates (NiV-Malaysia (NiV-M) and NiV-Bangladesh (NiV-B)), one *Pteropus* bat NiV isolate (NiV Cambodia (NiV-C)), and one equine HeV isolate. Viral entry into cells was quantified using recombinant vesicular stomatitis virus (VSV), which lacks envelope glycoprotein G, replaced by red fluorescent protein (rVSVΔG-RFP), pseudotyped with surface glycoproteins G and F from each Henipavirus isolate. Bat PCs, bat RSCs, and Vero cells were infected with pseudotyped rVSVΔG-RFP viruses at MOI of 0.01 PFU/cell and the percentage of infected cells was analyzed by flow cytometry 6 h later (Fig. 4a). In contrast to bat PCs, bat RSCs were susceptible to infection with all four tested viruses. Henipaviruses entered bat RSCs much more readily than bat PCs and, interestingly, more readily than Vero cells, which are commonly used to isolate and propagate *Henipavirus*. Next, we examined the transcription and replication kinetics of NiV by RT-qPCR and viral titration after infection of cells with virus at a low MOI (0.1 PFU/cell) (Fig. 4b, c). In bat RSCs, NiV RNA synthesis and production of viral particles increased by 4 log10 units between 0 h and 24 h post-infection; no increase was observed in bat PCs and all of the tested strains replicated much better in bat RSC than in bat PSC. Cytopathic effects and formation of giant multinucleated cells are hallmarks of NiV infection; both were readily visible in bat RSCs at 24 h post-infection (Fig. 4d) and developed further during the course of infection. These results demonstrate that unlike bat PCs, bat RSCs are highly permissive for *Henipavirus* infection, thereby suggesting novel opportunities for isolation of new bat viruses.

**Fig. 4.**
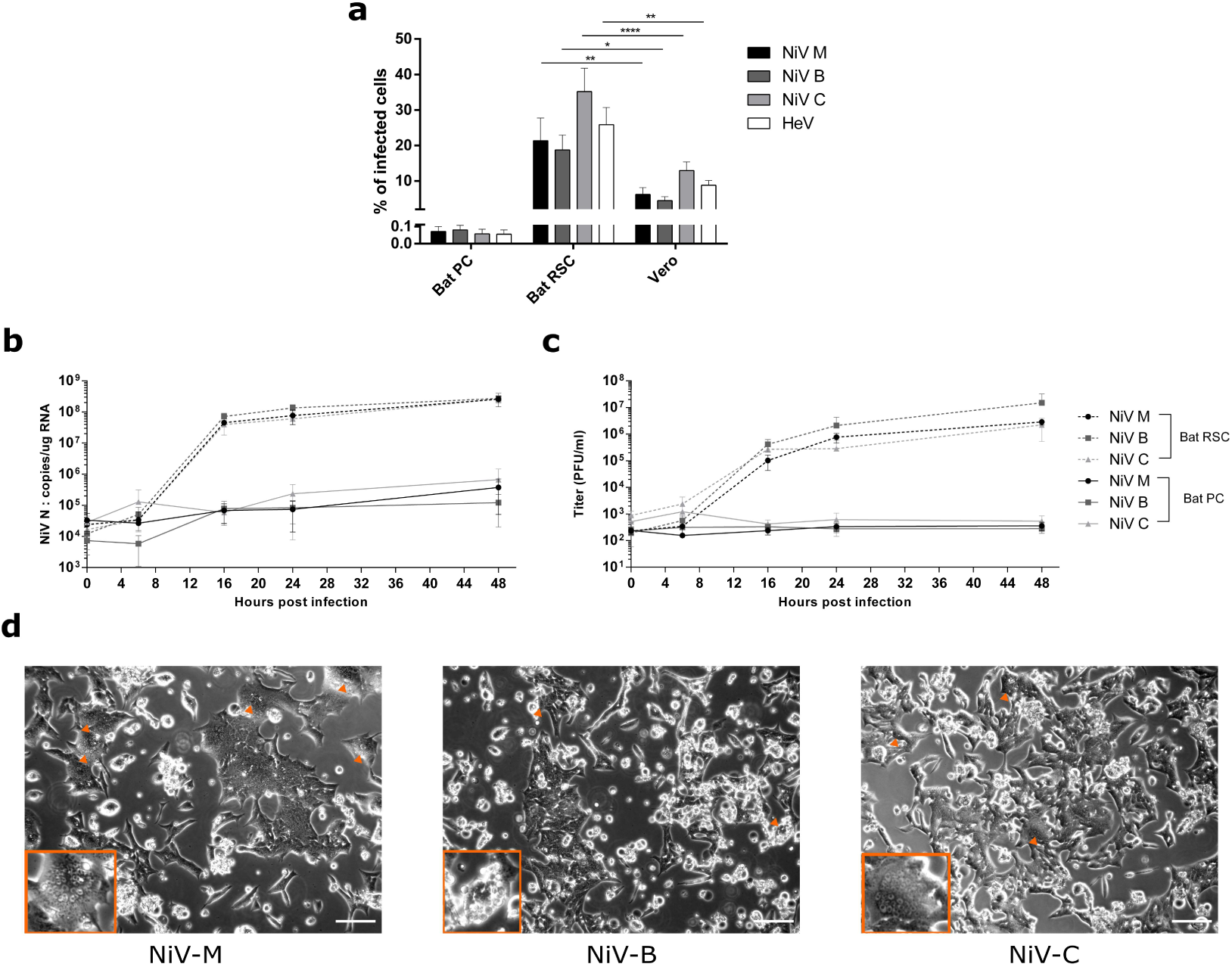
Susceptibility of *Pteropus* bat reprogrammed cells to Henipaviruses. (a) Entry of VSV pseudotyped with Henipavirus glycoproteins into bat PC, bat RSC, and Vero cells (MOI = 0.01 PFU/mL). Data were analyzed using One-Way ANOVA, followed by Tukey’s multiple comparisons test (**** P < 0.0001; ** P < 0.01; and * P < 0.05). (b–d) Bat PCs and bat RSCs were infected with the indicated viruses at a MOI of 0.1 PFU/mL. RT-qPCR analysis of Nipah virus infection kinetics (b) and release of virions into the supernatant (c). Bars represent the means ± SDs of three independent experiments. (d) Cell cytopathic effects observed under a light microscope at 24 h post-infection. Arrowheads show syncytia. Scale bar, 25 µm.

### Innate immune responses of bat RSCs

As innate immunity is the first line of defense against viral infection, we next analyzed innate responses of bat RSC and compared them with those of bat PCs. First, we performed a comparative analysis of ISG expression in non-stimulated bat PCs and RSCs at the global transcriptomic level. Taking advantage of a consensus list of ISGs^36^, we identified differential expression of 174 ISGs in the two cells types (Fig. 5a). Analysis identified two main sets of genes, one expressed at higher levels in bat RSCs (14 genes with a LFC > −2) and another expressed at higher levels in bat PCs (83 genes with a LFC > 2) (Supplementary Table 5). The second set contained a much higher number of genes, suggesting that reprogrammed cells expressed fewer ISGs and that these cells had weaker antiviral capacity. In particular, the interferon-regulated factor 1 (IRF1) gene network was down-regulated in bat RSCs (Fig. 5b, Supplementary Table 6). This specific intrinsic ISG signature may be one of the mechanisms underlying higher susceptibility of bat RSCs to infection by *Henipavirus*. Further functional approaches will help to identify which specific genes among those differentially expressed are implicated in bat antiviral responses.

**Fig. 5.**
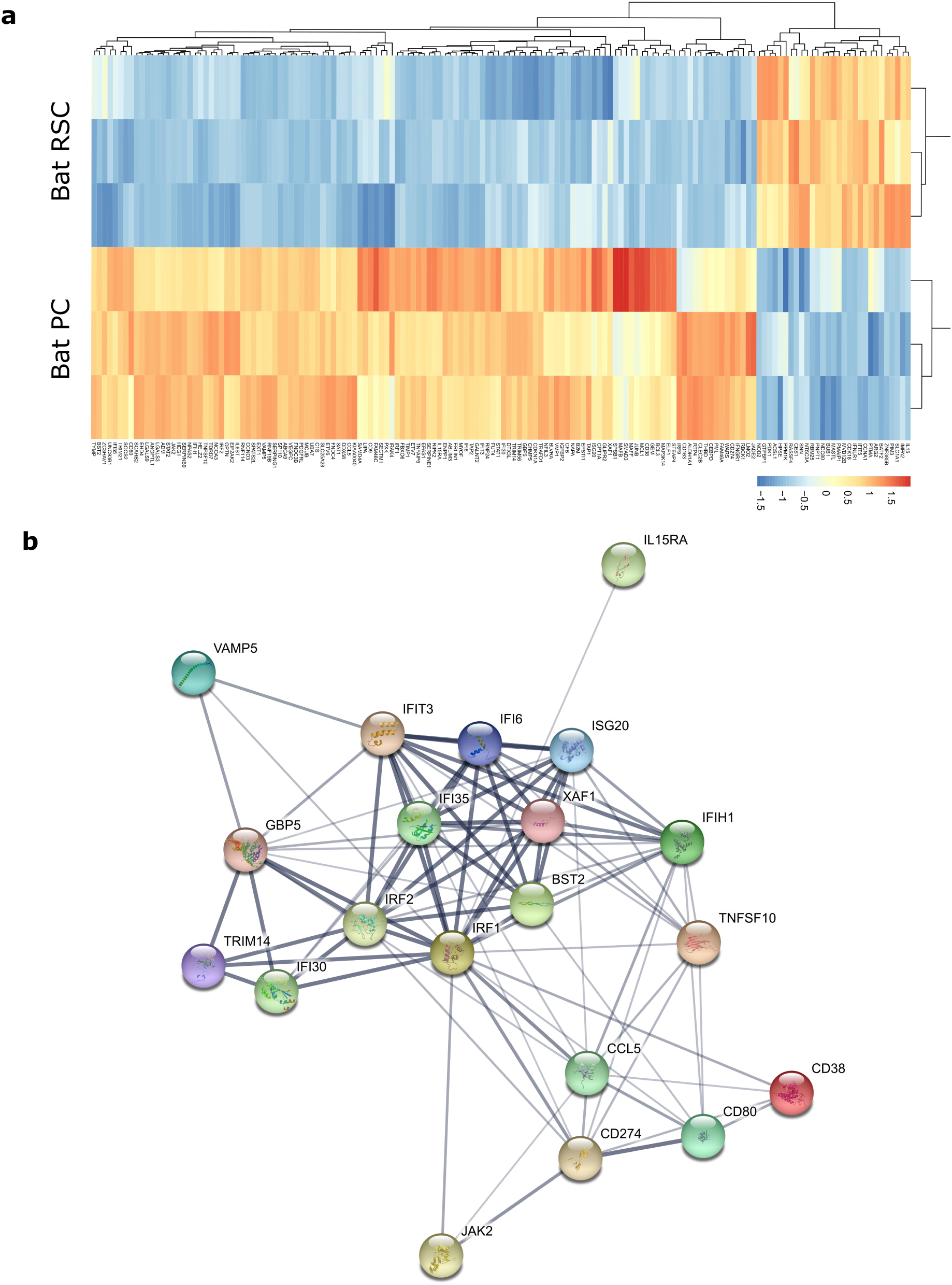
Bat RSCs exhibit an original ISG signature. (a) Heatmap showing the ISG signature in bat PCs and bat RSCs, illustrating the presence of two main set of genes: one is up-regulated in bat PCs and one (a larger set) is down-regulated in bat RSCs. (b) Cluster of ISGs centered on IRF1 and IRF2 following String analysis of ISGs down-regulated in bat RSCs.

### Bat RSC express IFN-I and ISGs following viral infection and respond to IFN-I stimulation

Finally, we compared innate immune response of bat RSCs and bat PCs in response to IFN-I. Initially, we analyzed the effects of polyI:C, a synthetic analogue of dsRNA, which binds and activates TLR3 when directly added to the cell culture^37^ and is used widely to mimic viral infection^38^. Only bat PCs produced both IFNα and IFNβ, as well as MX1, ISG56, and PKR, in response to increasing concentrations of polyI:C; no increase in ISGs expression was observed in stimulated bat RSCs (Fig. 6a). Next, we analyzed the effect of infection with a negative strand RNA virus, VSV, which stimulates the TLR3, RIG-I, and MDA5 pathways (Fig. 6b). In contrast to stimulation with polyI:C, VSV induced similar responses in both bat RSCs and bat PCs: VSV activated expression of IFNα, IFNβ and ISG56 expression, but not that of MX1 or PKR. Finally, stimulation of bat RSCs and bat PCs with IFN-I, known to activate the intracellular JAK-STAT pathway, induced similar responses in both cell types, characterized by IFN dose-dependent expression of MX1, ISG56, and PKR mRNA (Fig. 6c). These results suggest that bat RSCs produce and respond to IFN-I, similar to bat PCs; however, some intracellular signaling pathways, especially those related to the TLR3 pathway, seem to be activated differently.

**Fig. 6.**
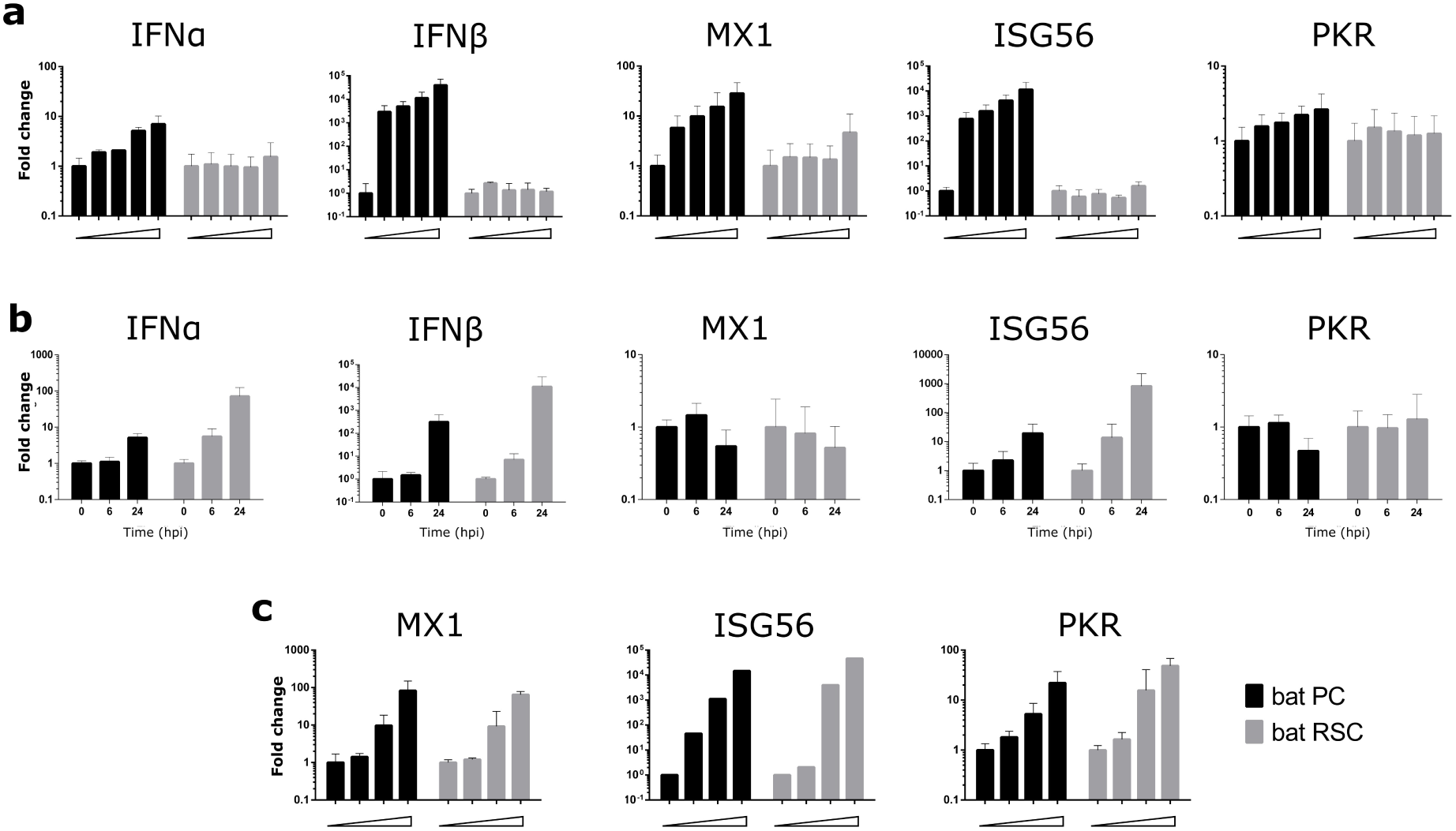
Innate immune responses of *Pteropu*s bat cells. (a) Dose-dependent induction of bat IFNα, IFNβ, Mx1, ISG56, and PKR transcripts in bat PCs and RSCs upon treatment with polyI:C, (b) VSV infection (MOI = 1 PFU/mL) and (c) IFN-u. Cells were stimulated 6 h with either polyI:C or IFN-γ, or infected with VSV for 6 or 24 h. Bars represent the means ± SDs of three independent experiments.

## Discussion

Efforts to combat deadly zoonotic infections caused by new emerging viruses are limited by lack of knowledge about reservoir host biology and the absence of adequate *in vitro* models for studying host-pathogen interactions^24^. Isolation of new viral strains is an important step and is (generally) attempted in established immortalized cell lines such as IFN-I-deficient simian Vero cells. Propagation of viruses in cell lines other than those of the natural host may lead to selection of virus sub-populations that replicate more efficiently in the chosen cell type, leading to accumulation of adaptive mutations that may prevent adequate assessment of host-pathogen interactions^23^. Therefore, further advancement in the field requires establishment of new bat cell models; indeed, development of novel *Pteropus* cell lines is particularly important for the field of *Henipavirus* infection research. The high permissiveness of bat RSCs reported herein (even higher than that of Vero cells routinely used for virus isolation and production) provides an important new tool for further studies. This cell model should bridge an existing gap in this field, facilitate optimization of viral discovery in bats^39^, and allow further research into bat biology at the cellular and molecular level.

In recent years, somatic reprogramming has become a powerful method for generating PSCs in species from which embryos or tissues are difficult to obtain. Using the original combination of ESSRB, CDX2 and c-MYC transcription factors, we obtained bat RPCs that exhibit stem cell-like properties. Unexpectedly, the OSKM reprograming factors OCT4, SOX2, KLF4, and c-MYC, which are used successfully in other mammalian species^26^, were inefficient in *Pteropus* cells, either alone or in combination with NANOG. This underlies the potential distinctiveness of bat cells with respect to this approach. Future somatic reprograming in other bat species will show whether the approach reported herein for *Pteropus* bat cells can be applied to other species. The reprogramming process presented in this report generated cells with rather unique stem cell characteristics. In addition to expression of PSC-specific antigens and epigenetic markers, these reprogramed bat cells express an original transcriptomic signature comprising numerous neural stem cell markers, as evidenced by GO terms and gene cluster analyses. As neural cells are one of the principal targets of Henipaviruses, this cell profile may favor infection, as observed in our study. In addition, as is currently the case for human stem cells^40^, further differentiation of reprogrammed bat stem cells into neural cell types relevant for viral infection may open new avenues for modeling of viral infectious diseases.

Recent studies of innate immune responses in human and murine stem cells reveal that mammalian ESCs and other types of PSCs do not have a functional IFN-based antiviral mechanism^41–43^; this is in contrast to avian ESCs that elicit antiviral responses similar to those of differentiated cells^44^. Attenuated innate immune responses by pluripotent human cells correlates with high expression of suppressor of cytokine signaling 1 (SOCS1)^45^. However, comparative analysis of RNAseq data in our study revealed low differential expression of SOCS1 in RSCs compared with that in PCs (LFC of −2.6), highlighting an additional distinction between bat and other mammalian cells. Absence of IFN-I expression in murine embryos is attributed to developmental regulation of IRF1 and IRF2 expression, which becomes functional after cell differentiation and is capable of regulating IFN-1 production^46^. Similarly, expression of IRF1 and IRF2 is lower in bat RSCs that in bat PCs (Supplementary Table 2), suggesting that another mechanism regulates IFN-I production by bat stem cells. Furthermore, previously observed constitutive expression of IRF7 in bats is thought to allow them to respond more rapidly to infection^47^. Although we did not find similar expression of IRF7 in our bat PCs and RSCs, we did (surprisingly) identify increased expression of IRF6 (Supplementary Table 2). While IRF6 is the only member of IRF family not linked to IFN control in mammals, it acts as a positive regulator of IFN expression in fish^48^ and inhibits production of the proinflammatory cytokine IL-1β by human cells^49^, suggesting that it may be involved in controlling innate immune responses and inflammation in bats; however, this needs to be further tested in functional studies.

Previous genome analysis of *Pteropus alecto* and *Myotis davidi* bats suggests that positive selection of components of the DNA damage checkpoint pathway is associated with changes in the overlap between this pathway and the innate immune system, indicating that evolutionary adaptations important for flight may have secondary effects on bat immunity^50^. Analysis of ISG expression in immortalized *P. alecto* bat cell lines reveals strong evolutionary conservation, although unstimulated bat cells express ISGs at higher level than their humans counterparts^51^. The profile of ISGs expressed in our unstimulated primary and reprogrammed bat cells is different from that expressed by immortalized cell lines, possibly due to absence of immortalization-activated programs, which may have affected ISG expression by bat cell lines in the previous study. Important differences in innate immune signatures between undifferentiated stem cells and their differentiated progeny have been described for several mammalian species; these differences lead to high resistance of stem cells to viral infection^36^. While we also observed clear differences between primary and reprogrammed bat cells (i.e., lower ISG expression in the latter), these unique cells are capable of both producing IFN-I and replicating viruses, including Henipaviruses; this is in sharp contrast to many other mammalian stem cells, but interestingly, similar to observations in avian stem cells^44^.

Bats possess peculiarities in terms of immune responses^22^, and our reprogrammed bat cells seemed to follow the same pattern. One of the first lines of defense against viral pathogens is the IFN response; indeed, contraction of the IFN-I locus and constitutive expression of IFNα was observed in *P. alecto* bats^52^. Interestingly, we did not find constitutive expression of IFN-I by bat stem cells or PC lines, although expression of IFNα and IFNβ was inducible in both cell types. In agreement with our results, a previous study did not observe constitutive expression of IFNα in cell lines generated from *R. aegyptus* bats^53^. Taken together, our data suggest that certain differences between in bat immune responses and those of other mammals are present at the stem cell level and may have been established during evolution, thereby shaping development of bat antiviral responses. Early exposure of pregnant bats to viral infections circulating in bat colonies may lead to development of disease tolerance in their progeny^54^ and the absence of a strong immune response to viral proteins; this may constitute a part of the antiviral bat defense strategy^55^. As suggested in a recent study of the *Rousettus* bat genome^53^, immune tolerance rather than a heightened antiviral defense strategy may support fitness, despite replication of pathogens, thereby allowing bats to efficiently control infection by viruses that are highly pathogenic to other mammalian species. In conclusion, we obtained reprogrammed bat cells presenting stem cell-like features and high permissiveness for *Henipavirus* infection. The particular innate immune signature of these cells underlines the distinctiveness of bat species and improves our understanding of the mechanisms underlying bat antiviral responses.

## Materials and methods

### *Pteropus* bat primary cell culture

Cell cultures of *Pteropus* bat flying fox (*Pteropus giganteus*, also known as *P. medius*) and *P. vampyrus* (*Yinpterochiroptera* suborder) were generated from samples collected in Tiergarten Schönbrunn (Vienna, Austria). Lung and trachea samples were obtained a female specimen found naturally dead, and wing-membrane skin biopsies were obtained during the regular veterinary checkup in accordance with national guidelines to minimize stress. Samples were washed with sterile PBS and transferred to Cryo-SFM freezing medium (PromoCell Bioscience, C-29910) on dry ice for shipment from the zoo. To obtain bat primary cell cultures (bat PCs), samples were dissected in a Petri dish and explants derived from the trachea (PTCs), lung (PLCs), and alary membrane (PACs) were cultured at 37°C/7% CO2 in parallel in gelatin-coated wells containing either fibroblast medium (FM) (DMEM/F12 supplemented with 10% fetal bovine serum (FBS), 2 mM L-glutamine, 1000 U/mL penicillin, 1000 U/mL streptomycin) or ESM1 medium (DMEM/F12 supplemented with 10% FBS, 2 mM L-glutamine, 1000 U/mL penicillin, 1000 U/ mL streptomycin, 1% non-essential amino acids (100X), 1 mM sodium pyruvate, 0.1 mM β-mercaptoethanol, 1 ng/mL IL-6, 1 ng/mL IL-6 receptor, 1 ng/mL Mouse Stem Cell Factor, 5 ng/mL insulin-like growth factor-1, and 1000 U/mL leukemia inhibitory factor); this culture protocol is used routinely for avian stem cells^56^. References to all reagents used for cell culture are listed in Supplementary Table 7.

### Reprogramming vectors

CDNAs encoding human POU5F1, SOX2, KLF4, NANOG, c-MYC, and CDX2, and mouse ESRRB were cloned, sequenced, and inserted into an inducible pPB transposon backbone. All constructs were generated using the NEBuilder® HiFi DNA Assembly Master Mix system (New England BioLabs, E2621) and sequenced to validate correct cDNA insertion. Viral stocks of the Sendai viral vector CytoTune2.0 kit, expressing the POU5F1/OCT4, SOX2, KLF4, and c-MYC human genes, were purchased from Invitrogen (A16517).

### Generation of reprogrammed *Pteropus* bat cells

Bat PCs were seeded in a 6-well plate and infected with viruses containing KLF4-OCT4-SOX2 and c-MYC (5 PFU/cell), and with a virus containing KLF4 (3 PFU/cell), according to the protocol provided by the virus supplier (CytoTune2.0). Cells were passaged 5 days after infection and seeded in two 55-cm^2^ flasks. The cells were cultured for 6 days in ESM1 or EPI medium (the medium was changed every 2 days). The EPI medium is serum-free and comprises 50% DMEM/F12 and 50% Neurobasal medium, supplemented with B-27 Supplement, N-2 Supplement, 2 mM L-Glutamine, 1000 U/mL penicillin, 1000 U/mL streptomycin, 1 mM β-mercaptoethanol, 5 ng/mL basic-Fibroblast Growth Factor (b-FGF), and 10 ng/mL human Activin A. In a second approach, bat PCs were modified by electroporation with different combinations of inducible transposons encoding OCT4, SOX2, KLF4, c-MYC, NANOG, ESRBB, and CDX2. Bat PC cells were dissociated and centrifuged at 1200 rpm (300 g) for 5 min at ambient temperature. The cell pellet was rinsed in PBS, centrifuged again, and 1.2×10^6^ cells recovered in 120 μL of R resuspension buffer (Neon, Life Technologies) containing 2 μg of pCAGPBase transposase-expressing vector and 4 μg of the reprogramming gene cocktail^31,57^. 100-μL of this cell-plasmid solution was electroporated (1500 V, 30 ms, 1 pulse) in a 100 μL tip (Neon, Life Technologies, MPK10096) using the Neon system (Life Technologies, MPK5000). After electroporation, the cells were cultured in 3 mL of FM medium in a 6-well plate. The medium from electroporated cells was replaced after 24 h and cells were selected using 5 μg/mL of puromycin and 200 μg/mL of neomycin, depending on the resistance genes carried by the plasmids present in the mixture. The medium containing selection reagents was changed every 2 days for at least 1 week. At the end of the selection process (between 8 and 15 days), the cells were dissociated by 0.05% trypsin-EDTA (Life) and 2×10^5^ cells were seeded into a well in a 6-well plate containing 3 mL of ESM1, EPI, or ESM2 medium supplemented with 2 µg/mL of doxycycline. The ESM2 medium comprised DMEM/F12 supplemented with 10% FBS, 2 mM L-glutamine, 1000 U/mL penicillin, 1000 U/ mL streptomycin, 1% non-essential amino acids (100X), 1 mM sodium pyruvate, 0.1 mM β-mercaptoethanol, 5 ng/mL b-FGF, and 10 ng/mL human Activin A. References for all reagents used for cell culture are listed in Supplementary Table 8.

### Characterization of *Pteropus* bat reprogrammed cells

Cells were examined by electron microscopy as described^34^. Exponentially growing cells were subjected to cell cycle analysis as described previously^58^. Reactivity of cells with antibodies specific for SSEA-1, SSEA-3, SSEA-4, or EMA-1 was tested as previously described^56^. The nuclear distribution of histone methylation marks was analyzed by immunofluorescence microscopy as described previously^34^. FastQ files were generated by RNAseq sequencing (Eurofins Genomics; https://www.eurofinsgenomics.eu/) of bat PTCs as a starting material for reprogramming, along with two reprogrammed cell populations, E2 and ES2, which were established by culture in EPI and ESM2 medium, respectively. For analysis, a reference index was created based on the *Pteropus vampyrus* annotation GCF_000151845.1_Pvam_2.0_rna.fna.gz from the NCBI database, which was updated on January 20, 2018. FastQ files were pre-processed using fastp v0.19.6 and then pseudoaligned using kallisto v0.8.1, with the following parameters: kallisto quant -i pVam2.0 --bias -b 100 -l -s -t 8^59,60^. DEGs were calculated from abundance.h5 files using standard parameters within DESeq2 v1.24.0^61^. An ISGs table^36^ was crossed with PTCvES DSGs, with an adjusted p-value lower < 0.05 using common gene symbols to generate an ISG heatmap. Raw data and gene tables are available through the GSE134585 datasets. Clustering and GO annotations were performed by String (https://string-db.org/) and Gene Ontology (http://geneontology.org/), using the human Gene symbols as references.

### Pseudotyped virus infection assay

Henipavirus pseudotyped particles were generated from rVSV-ΔG-RFP, a recombinant VSV in which the G protein envelope has been replaced with RFP (red fluorescent protein), as previously described (Gaudino, submitted). Briefly, the attachment glycoprotein and fusion proteins of NiV/Malaysia, NiV/Bangladesh, NiV/Cambodia, and HeV were cloned from RNA isolated from each virus and inserted into a pCAGGS plasmid vector. For each viral isolate, plasmid vectors coding the glycoproteins were transfected into BSR-T7 cells using the TransIT®-LT1 Transfection Reagent (Mirus Bio, MIR 2300). At 16 h post-transfection, cells were infected with rVSV-ΔG-RFP (0.3 PFU/cell) to produce a pseudotype VSV for each isolate. Supernatants were collected 24 h post-infection and concentrated by ultracentrifugation (28,000 rpm for 2 h at 4°C). Virus stocks were titrated on Vero cells. Pseudotyped viruses were named after the virus providing the surface glycoprotein. To evaluate viral entry into different bat cells, cells cultured in 24-well plates (80% confluent and adherent) were infected at an MOI of 0.01 PFU/cell for 1 h at 37°C. Then, virus-containing medium was removed and cells were washed once with PBS. Finally, fresh medium was added to the cells for 6 h at 37°C. The percentage of infected cells was assessed by measuring RFP fluorescence by flow cytometry (BD LSR Fortessa, BD Biosciences).

### Henipavirus infection and titration

NiV-Malaysia virus (isolate UMMC1; GenBank AY029767), NiV-Bangladesh virus (isolate SPB200401066, GenBank AY988601), and NiV Cambodia virus (isolate NiV/KHM/CSUR381, GenBank MK801755) were generated and titrated on Vero E6 cells at the INSERM Jean Mérieux BSL-4 laboratory in Lyon, France. For each virus, bat cells were plated in 12-well plates and upon reaching 80% of confluence were infected for 1 h at 37°C with virus at 0.1 PFU/cell. Next, virus-containing media were removed and cells were washed once with PBS. Finally, fresh medium was added to cells, followed by incubation at 37°C for 0, 6, 16, 24, or 48 h. Cell morphology was observed under a Zeiss Axiovert 100M microscope and photos were taken at 24 h post-infection using ImageJ software. For each time point, infected cell lysates were prepared using RLT buffer (Qiagen, 79216) prior to RT-qPCR analysis according to a validated BSL-4 procedure. Supernatants were collected and kept at −80°C until titration in a plaque assay on Vero E6 cells. Infections and titrations were performed in a BSL-4 facility at Jean Mérieux (Lyon, France).

### Induction of innate immune responses

Bat cells were stimulated with a synthetic analogue of dsRNA, polyinosinic-polycytidylic acid, polyI:C (Invitrogen), VSV Indiana strain, or universal IFN-type (a recombinant human IFN-alpha protein hybrid; Pbl Assay Science, 11200-2). Bat cells seeded in 12-well plates at 80% confluence were treated for 6 h at 37°C with either poly IC (0–1000 ng/mL) or universal IFN-type I (0–1000 U/mL). Alternatively, cells were infected with VSV (MOI = 1) for 1 h at 37°C. After viral attachment, cells were washed once with PBS and fresh medium was added. Cells were incubated for 0, 6, or 24 h at 37°C. Infected cell lysates were prepared for RT-qPCR analysis.

### RNA extraction and RT-qPCR analysis

At the indicated time points, lysates prepared from infected and stimulated cells were collected and RNA extracted using a NucleoSpin RNA kit (Macherey-Nagel, 740955). The yield and purity of the extracted RNA was assessed using a spectrophotometer (DS-11 FX; Denovix). Extracted mRNA was reverse-transcribed using SuperScript™ III Reverse Transcriptase (Invitrogen, 18080). Real time PCR was performed using Platinum SYBR green qPCR SuperMix-UDG kit with ROX reference dye (Thermo Fisher Scientific, 11733038) and a StepOne plus PCR system (Applied Biosystems). Data were analyzed using StepOne v2.3 software (Applied Biosystems) and calculations were done using the 2^ΔΔCT^ method. Expression was normalized to that of glyceraldehyde 3-phosphate dehydrogenase (GAPDH), as described previously^62^. The primers used are listed in Supplementary Table 8.

### Statistical analysis

All analyses were performed using GraphPad Prism 8 (GraphPad Software). Data were analyzed using analysis of variance two-way ANOVA, followed by Bonferroni or Tukey’s multiple comparisons test. P values < 0.05 were deemed significant.

## Supporting information

Spplemental Table 5

## Acknowledgements

The authors thank Drs. Doris Preininger, Anton Weissenbacher, and Tiergarten Schönbrunn (Vienna, Austria) for *Pteropus* samples. We also thank Drs F. Enchery, K. Dhondt, J. Fouret, and C. Legras-Lachuer for help in initiating and realizing this work.

## Supplementary data

**Figure S1.**
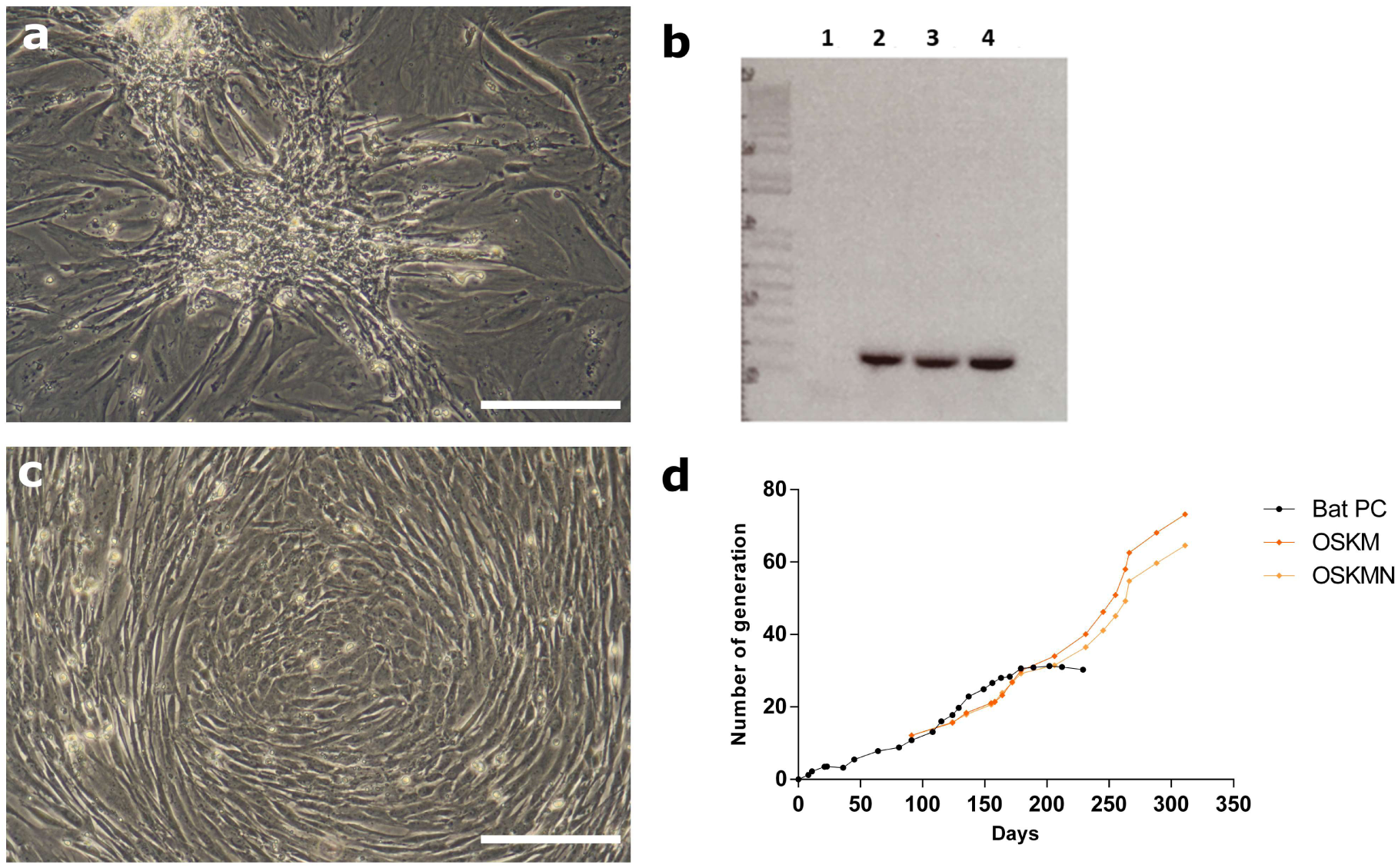
Attempts to obtain *Pteropus* bat reprogrammed cells using various delivery systems. (a) The classical OSKM combination was introduced into bat primary cells (PCs) using the non-integrative Sendai virus and cells were observed under a light microscope (scale bar, 200 μm). (b) PCR-based detection of Sendai virus was performed as recommended by the supplier: 1, bat PCs; 2, bat PCs + Sendai; 3, bat PCs + Sendai + NANOG; 4, Control reprogrammed stem cells. (c) Bat PCs were modified by electroporation of inducible transposons encoding OCT4, SOX2, KLF4, and c-MYC. Scale bar, 200 μm. (d) Comparative growth curves of bat PCs modified by electroporation of OSKM and OSKMN.

**Table S1 |** Pluripotent-associated genes identified in humans and *P. vamp*yrus bats.

**Figure 2e |** Figure 2e higher resolution of FACS analysis

**Table S2** | List of genes expressed by bat PCs (PTCs) and reprogrammed cells cultured in either EPI ECM_E2) or ESM (ECM_ES2) medium after RNAseq analysis

**Table S3 |**List of gene clusters identified after String analysis.

**Table S4 |** Functional enrichment of DEGs identified by RNAseq analysis.

**Table S5 |** List of ISG genes expressed by either bat PC or RSCs.

**Table S6 |** List of ISG genes clustered after String analysis.

**Table S7 |** List of reagents used for cell culture.

**Table S8 |** List of primers used in the study.

**Table S1.** Comparison of protein sequences of pluripotent genes between human and *Pteropus vampyrus* bat.

| **Gene** | ***Homo Sapiens***<br>**protein ID** | ***Pteropus vampyrus***<br>**protein ID** | **% alignment (proteins) between *Homo Sapiens* and *Pteropus vampyrus* species** |
| --- | --- | --- | --- |
| OCT4 | NP_002692.2 | XP_011382211.1 | 90.9 |
| SOX2 | NP_003097.1 | XP_011381167.1 | 99.1 |
| KLF4 | NP_001300981.1 | XP_011363252.1 | 87.3 |
| C-MYC | NP_002458.2 | XP_011359121.1 | 92.0 |
| NANOG | NP_079141.2 | XP_011364079.1 | 69.9 |
| CDX2 | NP_001256.3 | XP_011358539.1 | 91.0 |
| ESRRB | NP_004443.3 | XP_011355347.1 | 98.8 |

